# Tissue remodeling and cell signaling underpin changes in tumor microenvironment heterogeneity in glioma oncogenesis

**DOI:** 10.1101/2021.12.05.471299

**Authors:** Marija Dinevska, Samuel S. Widodo, Liam Furst, Lucero Cuzcano, Yitong Fang, Stefano Mangiola, Paul J. Neeson, Phillip K. Darcy, Robert G. Ramsay, Fabienne MacKay, Stanley S. Stylli, Theo Mantamadiotis

**Author notes:** Equal contribution (Marija Dinevska, Samuel S. Widodo). Correspondence should be addressed to T.M. current address: QIMR Berghofer Medical Research Institute, Brisbane. Australia.

## Abstract

Brain tumor cells thrive by adapting to the signals in their microenvironment. Understanding how the tumor microenvironment evolves during disease progression is crucial to deciphering the mechanisms underlying the functional behavior of cancer cells. To adapt, cancer cells activate signaling and transcriptional programs and migrate to establish micro-niches, in response to signals from neighboring cells and non-cellular stromal factors. Using multiple tissue analysis approaches to identify and measure immune cell infiltration and extracellular matrix deposition in brain tumors, we show that low-grade glioma is largely devoid of infiltrating immune cells and extracellular matrix proteins, while high-grade glioma exhibits abundant immune cell infiltration and activation, as well as extensive collagen deposition. Spatial analysis shows that most T-cells are sequestered in perivascular nests, but macrophages penetrate deep into tumor cell rich regions. High-grade gliomas exhibit heterogeneous PI3K and MAPK signaling, which correlates with distinct pathological hallmarks, including tumor angiogenesis, tumor cell density and extracellular matrix deposition. Our results also provide compelling evidence that tissue remodeling is an important element in glioma progression, and that targeting the extracellular matrix will be critical to improving GBM therapy.

## Introduction

Despite developments in targeted therapies and immunotherapy delivering success for some cancer types, primary malignant brain cancer remains incurable. In 2018, central nervous system cancers constituted 1.7% of all cancers, with an estimated 297,000 new cases worldwide, but account for 3.1% of cancer deaths (Wild, Weiderpass, and Stewart 2020). Gliomas account for approximately 70% of primary brain tumors and are divided into four pathological grades, encompassing low-grade glioma (LGG) and high-grade glioma (HGG). LGG typically progresses to HGG, with limited understanding of the cellular and molecular mechanisms involved in this transition. Grade IV HGG, known as glioblastoma (GBM) is the most aggressive and most common primary brain tumor in adults (Louis et al. 2016) with a dismal survival rate of approximately 15 months following diagnosis and standard therapy (Stupp et al. 2005). High-throughput gene expression analysis identified four distinct GBM subtypes and revealed underlying biological properties associated with the subtypes (Phillips et al. 2006), but little clinical benefit has accompanied this leap in scientific understanding. Large tumor cohort LGG and HGG next-generation DNA- and RNA-sequencing and GBM single cell sequencing investigations have revealed several layers of cellular and genetic complexity of the LGG and HGG microenvironment (Cancer Genome Atlas Research Network 2008; Brennan et al. 2013; Darmanis et al. 2017; Couturier et al. 2020) and the cellular origins of glioma (Castellan et al. 2021; Richards et al. 2021). The glioma tumor microenvironment harbors many cell types, with most investigations focusing on immune cell infiltration (Rahman et al. 2018; D. Yan et al. 2017; Andersen et al. 2021).

Of the non-cellular components of the tumor microenvironment, the extracellular matrix proteins (ECM) have been investigated in breast and lung cancer, and are implicated in several pro-oncogenic hallmarks, including tissue stiffness, tumor cell invasion and growth factor sequestration and release (García-Mendoza et al. 2016; Dinca et al. 2021; Barcus et al. 2021). Although the brain is deficient in inflammatory and immune cells, compared to other organs, pathological insults can trigger immune infiltration and activation, with associated pathological responses, including immune cell infiltration and tissue remodeling. Elevated expression of several ECM proteins, including collagen I, collagen IV, laminin and fibronectin have been identified in brain tumor tissue (Rutka et al. 1987). Recent work suggests that elevated collagen IV mRNA expression correlates with poor glioma patient survival, and collagen IV regulates glioma cell proliferation and migration (Wang et al. 2021). With the emerging view that tumors and wound healing share cellular and molecular mechanisms, including sustained proliferative signaling, activation of invasion, angiogenesis and inflammation (MacCarthy-Morrogh and Martin 2020);

ECM protein expression in glioma suggests that brain tumors share similar mechanisms to wound-healing. However, an understanding of the cellular and non-cellular components and their functional relationship in the glioma tumor microenvironment, remain elusive. Here we integrated multispectral immunohistochemistry, histopathological staining and spatial analysis to identify tumor-associated immune cells and their spatial relationship to key histopathological hallmarks in the glioma tumor microenvironment. By mapping the ECM-stromal architecture across glioma grades, we also identify the relationship between immune cell infiltration and tissue remodeling in glioma carcinogenesis.

## Results

### Early myeloid and late T-cell tumor infiltration

Resident microglia and infiltrating bone marrow-derived macrophages comprise the bulk of immune cells in brain tumor tissue (Quail and Joyce 2017). Tumor associated macrophages (TAMs) are considered important regulators of oncogenesis, exhibiting pro-tumor and anti-tumor functional states, and are increasingly viewed as important immunotherapeutic targets or cell-based therapies (Kowal, Kornete, and Joyce 2019; Quail and Joyce 2017). Using antibody panels to determine cell identity by multiplex immunohistochemistry (mIHC) (Table S1 & Figure S1), we measured immune cell composition and cell density in human non-tumor (‘normal adjacent to tumor’) brain tissue, LGG (grades I & II) and HGG tissues (grades III & IV) tissue (Figure 1a). In non-tumor brain tissue, microglia comprised 2.35% of all cells, at a density of 31 cells/mm^2^ (Figure 1 & Table S2), which is similar to previous reports (Dos Santos et al. 2020). The proportion of microglial cells was similar across all glioma tissue grades (Figure 1b). Macrophages were the most abundant infiltrating immune cell, comprising about 5% of all cells, with a median cell density of between 50 and 140 cells/mm^2^, across all tumor grades. T-cells were the least abundant immune cell type across all grades, with the highest cell number in grade IV glioma (GBM). Dendritic cells comprised about 1% of all cells across all tumor grades, with the largest proportion (1.7%) in grade I glioma (Figure 1b).

**Figure 1.**
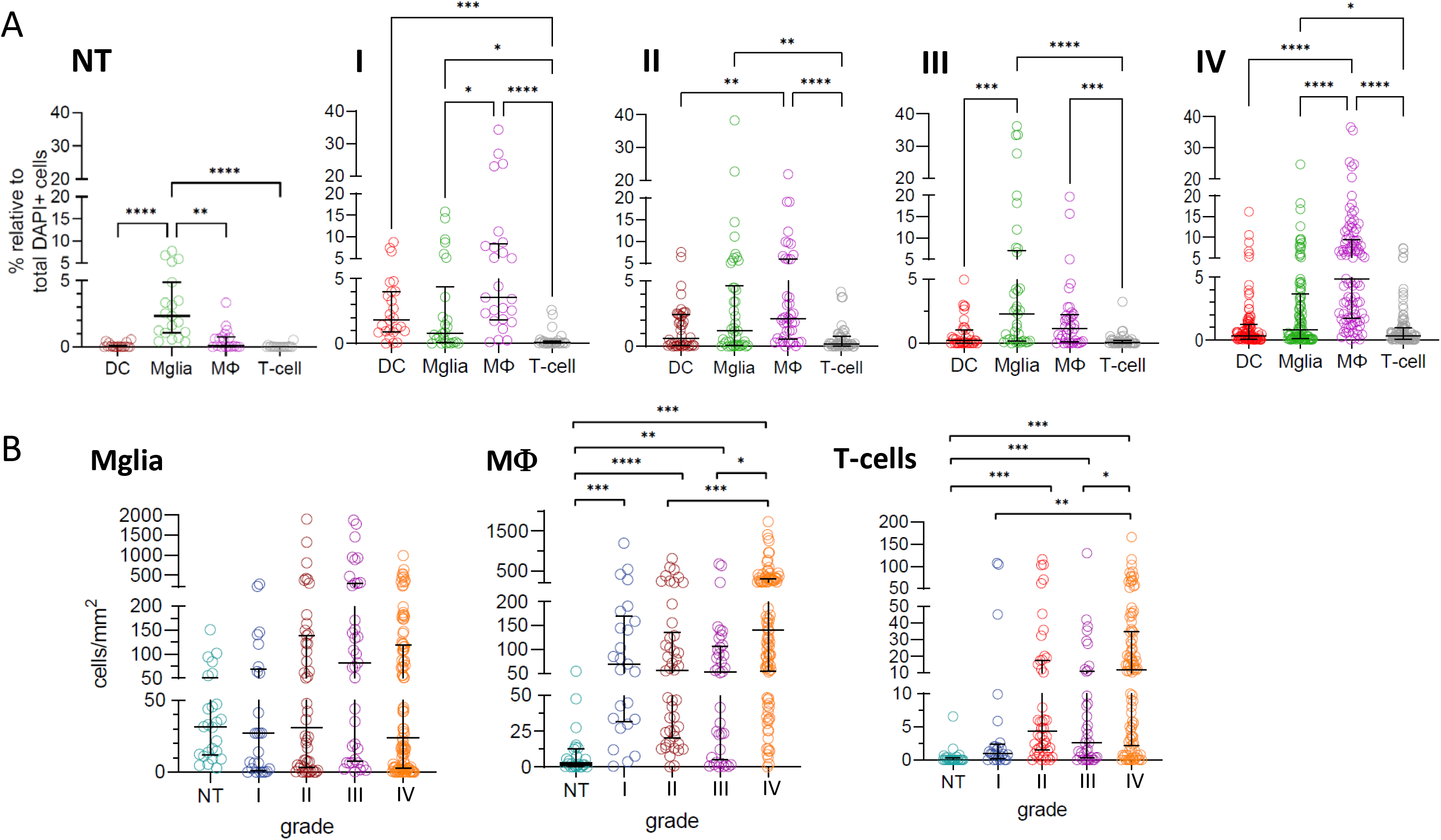
Grade-dependent immune cell infiltration, dominated by macrophages, correlates with tumor grade. **(A)** Immune cell composition and density in glioma grades I-IV and in non-tumor (NT) brain tissue were determined by mIHC image Halo analysis, using antibodies recognizing CD11c (dendritic cell; DC), TMEM119 (microglia; Mglia), CD68 (macrophage; MФ) and CD3 (T-cell). Immune cell composition is represented as the percentage of all (DAPI+) cells. **(B)**, Cell density is represented as the number of cells per area (mm^2^) analyzed. The data was derived by analyzing 182 glioma tissues (NT brain, *n*=19; grade I, *n*=22; grade II, *n*=42; grade III, *n*=34; grade IV, *n*=65). Statistical significance was determined using a Kruskal-Wallis test, followed by a Dunn’s test for multiple comparisons (α=0.05) and significance is represented by *(p < 0.05), **(p < 0.01), ***(p <0.001) and ****(p <0.0001).

Next, we investigated T-cell subsets and showed that in GBM, CD8+ T-cells were the most abundant cell type, constituting about 30% of all tumor infiltrating T-cells, followed by CD4+ T-cells, representing about 7.5% of all tumor infiltrating T-cells (Figure 2). This was reflected in the cell density of each T-cell subset (Figure 2B). Notably, FOXP3+ regulatory T-cells (Treg) cells were rarely detected and constituted less than 0.5% of T-cells. PD-1+ CD4+ and PD-1 CD8+ T-cells were also rare, present at a median density of less than 1 cell/mm^2^ (Figure 2B). To independently assess immune cell composition, we performed computational deconvolution analysis of LGG (Cancer Genome Atlas Research Network et al. 2015), GBM (Brennan et al. 2013) and non-tumor brain (Miller et al. 2014) gene expression data using CIBERSORT (Newman et al. 2015) and observed a similar immune cell profile to that measured in tissue (Figure S2).

**Figure 2.**
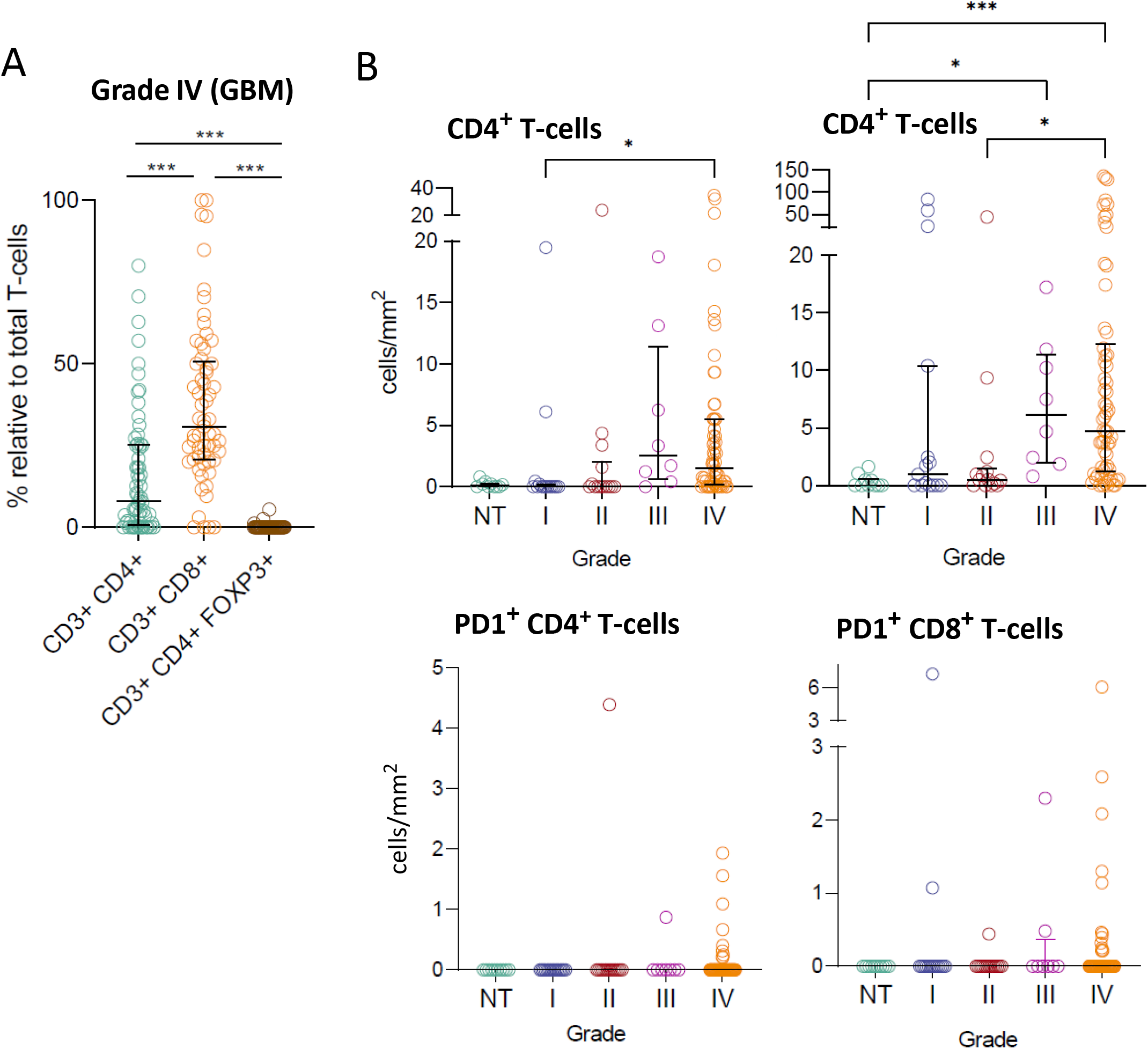
Most glioma tumor infiltrating T-cells are cytotoxic CD8+ and T-helper CD4+ cells, and immunosuppressive Treg and PD1+ T-cells are rare. **(A)** T-cell subset composition in grade IV (GBM) were identified using mIHC using CD3, CD8, CD4, FOXP3 and PD-1 antibodies and the data are represented as the proportion of cells relative to total T-cells (CD3+ DAPI+ cells). (B) T-cell density (cells/mm^2^) in glioma grades I-IV. The data was derived by analyzing 109 patient glioma tissues (non-tumor brain, *n*=10; grade I, *n*=15; grade II, *n*=14; grade III, *n*=8; grade IV, *n*=62). Statistical significance was determined using a Kruskal-Wallis test, followed by a Dunn’s test for multiple comparisons (α=0.05), significance is represented by *(p<0.05), **(p<0.01), ***(p<0.001) and ****(p<0.0001).

Computational deconvolution transcriptomic analysis showed that anti-inflammatory macrophages represented the largest proportion of immune cells in LGG (28%) and GBM (33%); CD4 T-cells constituted 20% of all immune cells in LGG, and 23% in GBM, respectively, and CD8 T-cells represented about 2% of all immune cells. Additional immune cell type gene signatures predicted that other cells were present at low abundance. Non-tumor brain cell transcriptomic data predicted the presence of low levels of immune cells, noting that the tissue used for gene expression analysis is mostly from pre- or perinatal origin, whereas most of the glioma expression data was derived from adult patient tissue.

### Immune cell activation correlates with tumor progression

To investigate immune cell activation in glioma tissue, we used CREB activation as a biomarker. CREB is a phosphorylation-dependent transcription factor expressed in all cells and is transiently phosphorylated by upstream serine-threonine kinases (Shaywitz and Greenberg 1999). CREB regulates multiple tumor and immune cell functions, including cancer cell proliferation, previously determined by co-expression with Ki-67 and proliferating cell nuclear antigen (PCNA) (P. M. Daniel, Filiz, Tymms, et al. 2018), T-cell proliferation (Baumann et al. 2004; Barton et al. 1996), metabolic suppression of tumor-infiltrating CD8+ T-cells (Mastelic-Gavillet et al. 2019), T-cell cytokine expression (Zhang et al. 2000), dendritic cell function (Ohl, Schippers, and Tenbrock 2018) and macrophage polarization (Luan et al. 2015). Using a phospho-specific CREB antibody in combination with immune cell type biomarkers, we observed a grade-dependent increase in the number of pCREB expressing immune cells, with T-cells exhibiting the greatest increase, followed by microglia, macrophages and dendritic cells (Figure 3A). The proportion of pCREB+ cells for specific immune cell types varied, with T-cells exhibiting the largest grade-dependent increase in pCREB+ cells. CD8+ pCREB+ T-cells were more abundant than CD4+ pCREB+ cells. pCREB+ dendritic cells, macrophages and T-cells were often detected in homogeneous cell clusters (Figure 3B).

**Figure 3.**
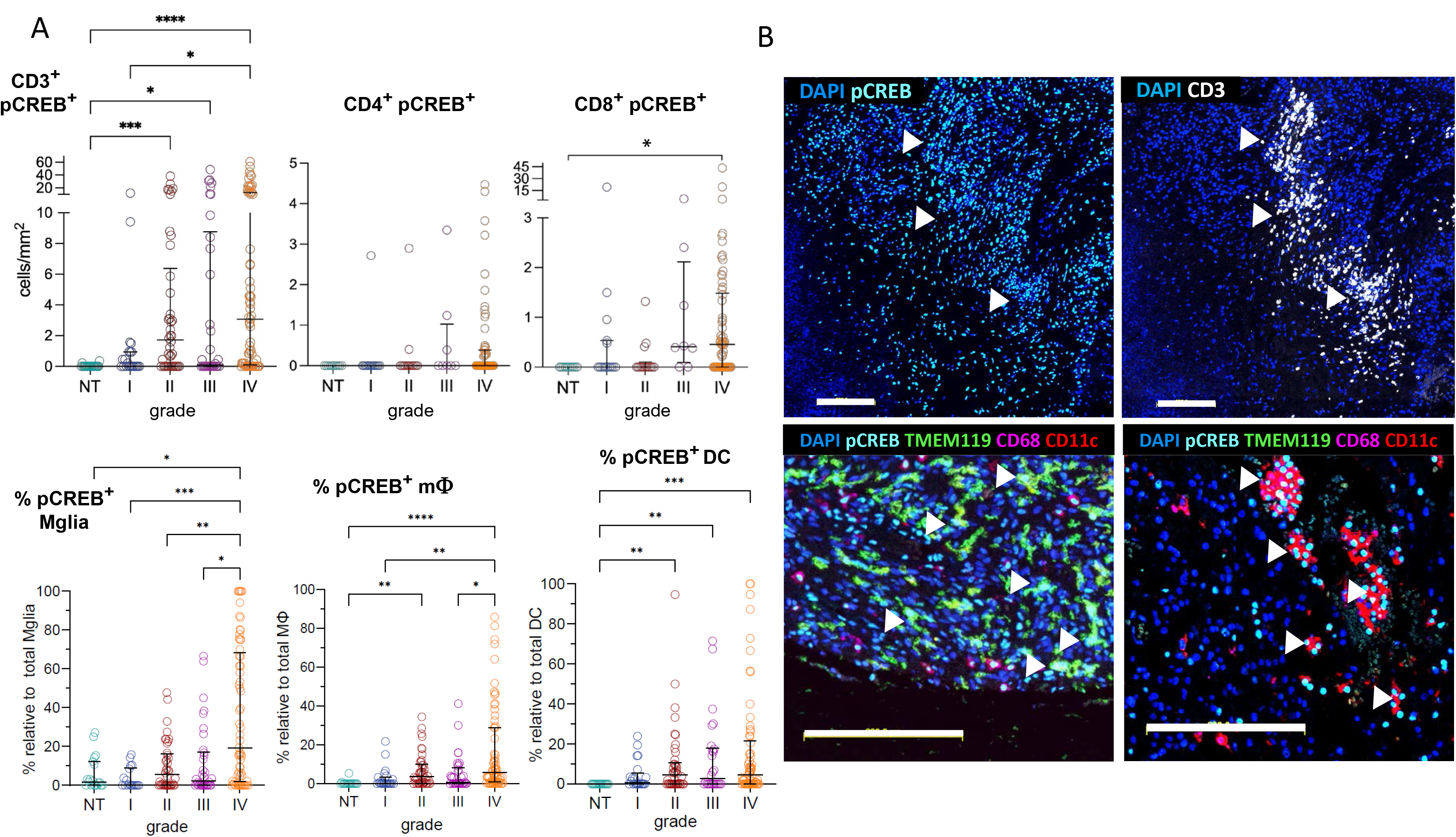
Grade dependent increase in immune cell activation. **(A)** The density (cells/mm^2^) of activated (pCREB+), total T-cells (CD3+) and CD4+ and DC8+ T-cell subsets increased with increasing grade. The proportion of activated dendritic cells, macrophages and microglia increased with grade. Statistical significance was determined using the Kruskal-Wallis test, followed by Dunn’s test for multiple comparisons (α=0.05), significance is represented by *(p<0.05), **(p<0.01), ***(p<0.001) and ****(< 0.0001). **(B)** The top two images are single channel images for pCREB+ and CD3+ and colocalization of these biomarkers, demonstrating activated T-cell clusters (white arrowheads). The white arrowheads in the bottom images indicate the presence of activated (pCREB+ TMEM119+) microglia (left image), and clustered activated pCREB+ CD68+ macrophages (right image). Scale bars are 200μm for all images.

### T-cell restriction within perivascular nests and stromal immune niches in GBM

We investigated T-cell distribution and localization in GBM, since grade IV tissue showed the highest T-cell density and T-cell clustering. Most T-cells were observed around blood vessels (Figure 4). Both CD4+ and CD8+ were localized to perivascular niches, with most cells in contact with the blood vessels or within 100μm of blood vessels (Figure 4), unlike macrophages which exhibited broad tissue distribution, both in perivascular niches and deep within tumor cell rich regions (Figure 4 B, C). Tertiary lymphoid structures were also observed around some blood vessels, which were composed of B-cells, T-cells and macrophages (Figure S3). As perivascular regions are typically rich in ECM proteins, we used Masson’s Trichrome staining to distinguish ECM-stroma and tumor cell rich regions. ECM protein deposition was observed at varying thickness around all blood vessels (Figure 4B and S3). To determine T-cell and macrophage distribution around blood vessels, mIHC was performed using an immune cell biomarker antibody panel in combination with a CD31 antibody to label endothelial cells, followed by Masson’s Trichrome staining. Masson’s Trichrome stained images were overlaid onto the mIHC immune cell data to identify ECM-stroma, tumor cell rich tissue regions and immune cell distribution.

**Figure 4.**
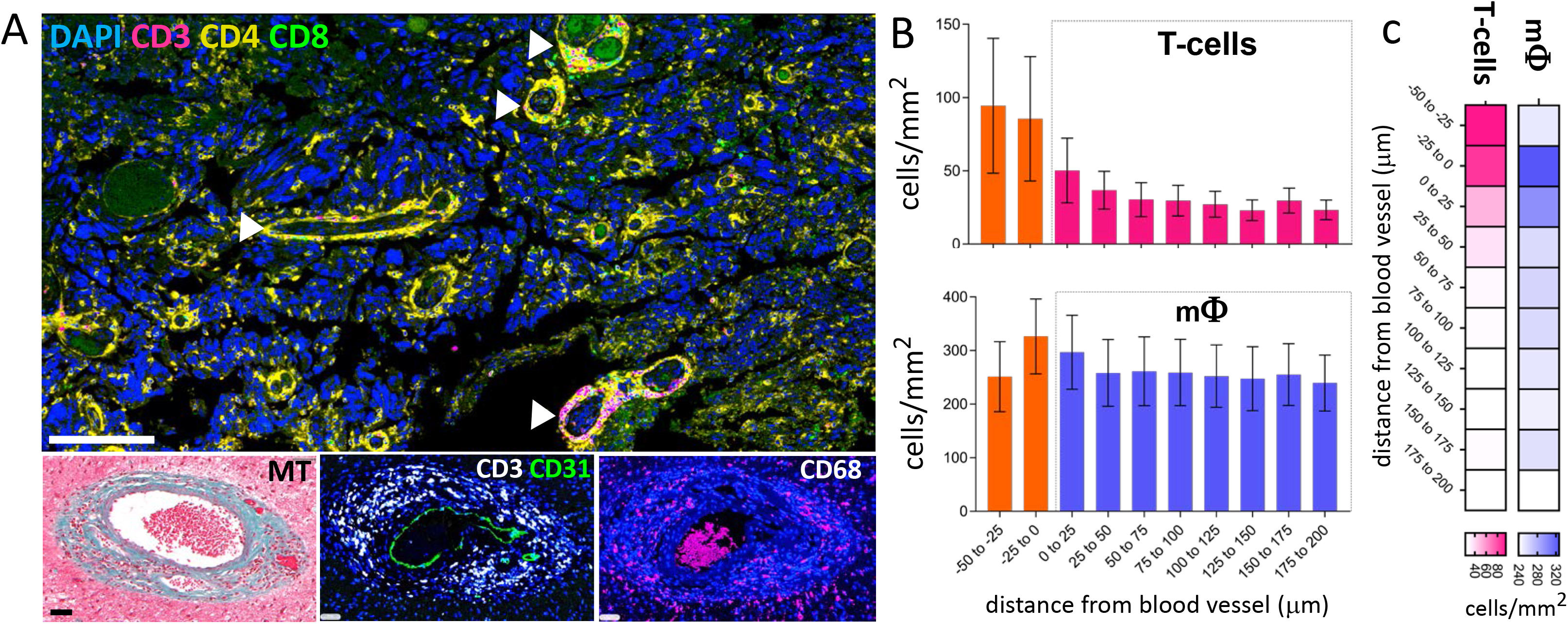
Perivascular immune cell niches. **(A)** mIHC demonstrates CD4+ and CD8+ T-cell clustering around perivascular niches in a GBM (upper image). T-cells (CD3+) and macrophages (CD68+) localized around endothelial cells (CD31+) and within a perivascular ECM-rich zone, indicated by the adjacent Masson’s Trichrome (MT) stained section. Scale bars are 100μm for the top, and 50μm for the bottom images. **(B)** T-cell (*n*=34) and macrophage (*n*=28) localization around the perivascular area and the infiltration into the tumor expressed as the distance from blood vessels in μm, was determined by overlaying mIHC and MT staining and spatial infiltration analysis using Halo software (see Methods). Error bars are +/- SEM. **c**, Relative immune cell density and tumor infiltration distance is depicted as a heatmap.

Thus, regional annotation combined with cell-specific spatial proximity analysis allowed a comparison of cellularity between ECM-stromal and tumor cell rich regions. Analysis confirmed the visual observation that T-cells were restricted to perivascular nests, but macrophages were broadly distributed in the ECM-stroma and in tumor cell rich tissue regions (Figure 4C). T-cells were present at a density of between 80 and 100 cells/mm^2^ within or directly in contact with blood vessels, and at a density of 50 cells/mm^2^ within 25μm of blood vessels. Beyond 25μm of the vascular tissue, T-cell density averaged between 10 and 20 cells/mm^2^ (Figure 4B). Macrophages were present at between 250 and 320 cells/mm^2^ and localized to perivascular nests and regions away from blood vessels, deep into tumor cell rich areas (Figure 4B).

### Extensive collagen deposition and tissue remodeling in GBM

Long-standing studies have reported the presence of stromal tissue in GBM (Perria and Sacchi 1950), but the stromal component has not been well characterized. To determine the extent of ECM deposition across glioma grades, tissues were stained with Masson’s Trichrome stain. LGG (grades I and II) and grade III tissue exhibited low ECM-stroma abundance, with less than 5% of the tissue area staining positive for collagen/ECM (Figure 5). GBM tissue showed variable ECM deposition, ranging from no visible ECM, to almost complete ECM coverage in the tissue cores examined. Examination of 20 larger GBM tissue samples (>10mm^2^) showed that extensive ECM deposition was common in GBM (Figure 5A). ECM deposition was higher in recurrent GBM tissue compared to primary GBM (Figure 5B), and higher in tissue from male patients over 40 years old (Figure 5B).

**Figure 5.**
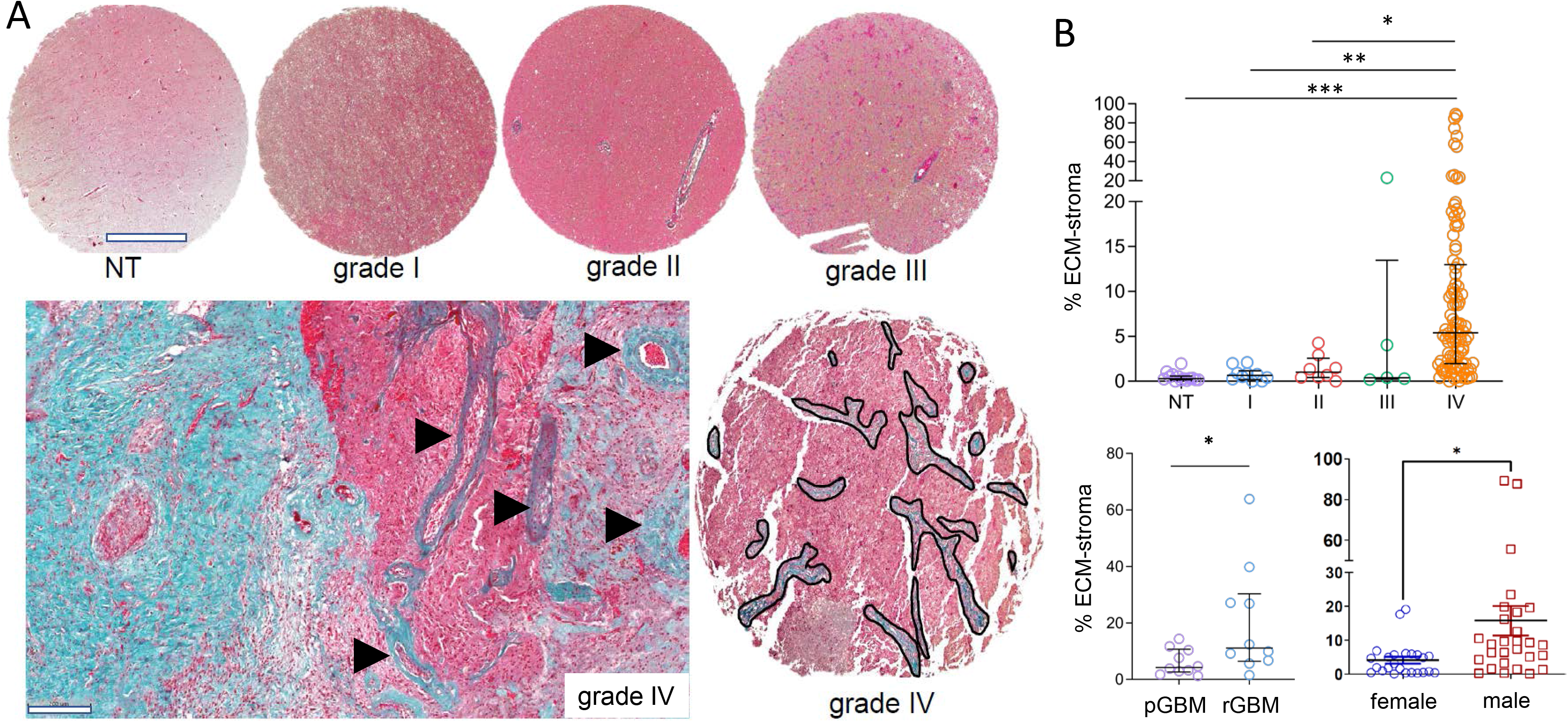
Grade-dependent ECM-stroma deposition. **(A)** Representative Masson’s Trichrome staining of adjacent non-tumor (*n*=14), grade I (*n*=10), grade II (*n*=8), grade III (*n*=5) and grade IV (*n*=100) tumor cores demonstrating increased presence of collagen with tumor grade (annotations indicating collagen). Arrowheads indicate perivascular collagen deposition. Scale bars are 500μm for the cores, and 200μm for the bottom image. **(B)** Percentage of ECM-stroma in the non-tumor and tumor cores (grade I-IV), and in primary (*n*=10) and recurrent (*n*=10). GBM whole tissue was determined using the Tissue Classifier Add-on in Halo™ (see Methods). The percentage of ECM-stroma in female and male patients over 40 years old was also compared (n=24 female; 28 male). Error bars are +/- IQR. Statistical significance was determined by Kruskal-Wallis test, followed by Dunn’s test for multiple comparisons (α=0.05) for comparison of ECM-stroma between tumor grades or by a two-tailed unpaired Student’s t-test for comparisons between primary/recurrent and male/female. Significance is represented by *(p<0.05), **(p<0.01), ***(p<0.001) and ****(p<0.0001).

To investigate differences in cell composition between stroma and tumor cell rich regions, we measured immune cell density and endothelial cells in ECM-stroma and tumor cell rich regions. The most abundant immune cell type, macrophages, were present at a small but significantly higher cell density in the ECM-stroma, compared to tumor cell rich regions (Figure 6B). CD31+ endothelial cells evident as blood vessels were enriched in the ECM-stroma and largely absent in tumor rich (CD44+) regions (Figure 7A and S4).

**Figure 6.**
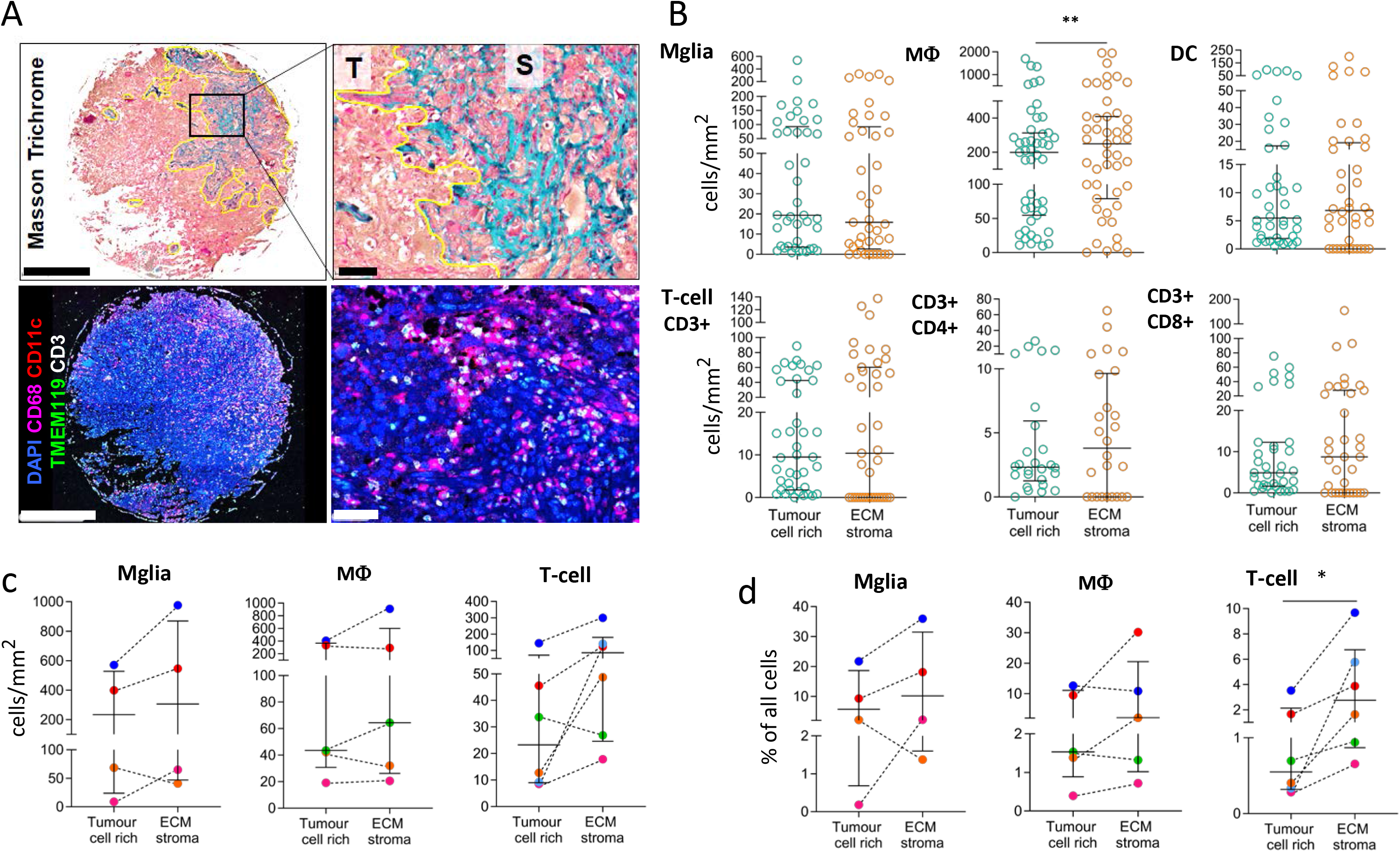
Tumor infiltrating immune cells are enriched in the ECM-stroma. **(A)** Representative Masson’s Trichrome staining and mIHC stained overlay for CD68, CD11c, TMEM119 and CD3 showing a tumor cell dense region (T) and ECM-stroma (S). Scale bars are 500μm for images on the left, and 50μm for images on the right. **(B)** Cell density of microglia (Mglia) (*n*=41), macrophages (MΦ) (*n*=47), dendritic cells (DC) (*n*=40), CD3+ (*n*=39), CD3+CD4+ (*n*=26) and CD3+CD8+ T (*n*=35) cells is shown as the number of cells per tumor cell rich and ECM-stroma area analyzed (mm^2^) within tumor cores. **(C)** Cell densities of microglia (Mglia) (*n*=4), macrophages (MΦ) (*n*=5), and T cells (*n*=6) are shown as the number of cells per tumor cell rich, and ECM-stroma area analyzed (mm^2^) within the whole tumor sections. **(D)** Microglia (Mglia) (*n*=4), macrophages (MΦ) (*n*=5), and T cells (*n*=6) are represented as the proportion of cells relative to total nucleated cells (DAPI+ cells). Error bars are +/- IQR. Statistical significance was determined by Wilcoxon matched-pairs signed rank test and significance is represented by *(p<0.05), **(p<0.01), ***(p<0.001) and ****(p<0.0001).

**Figure 7.**
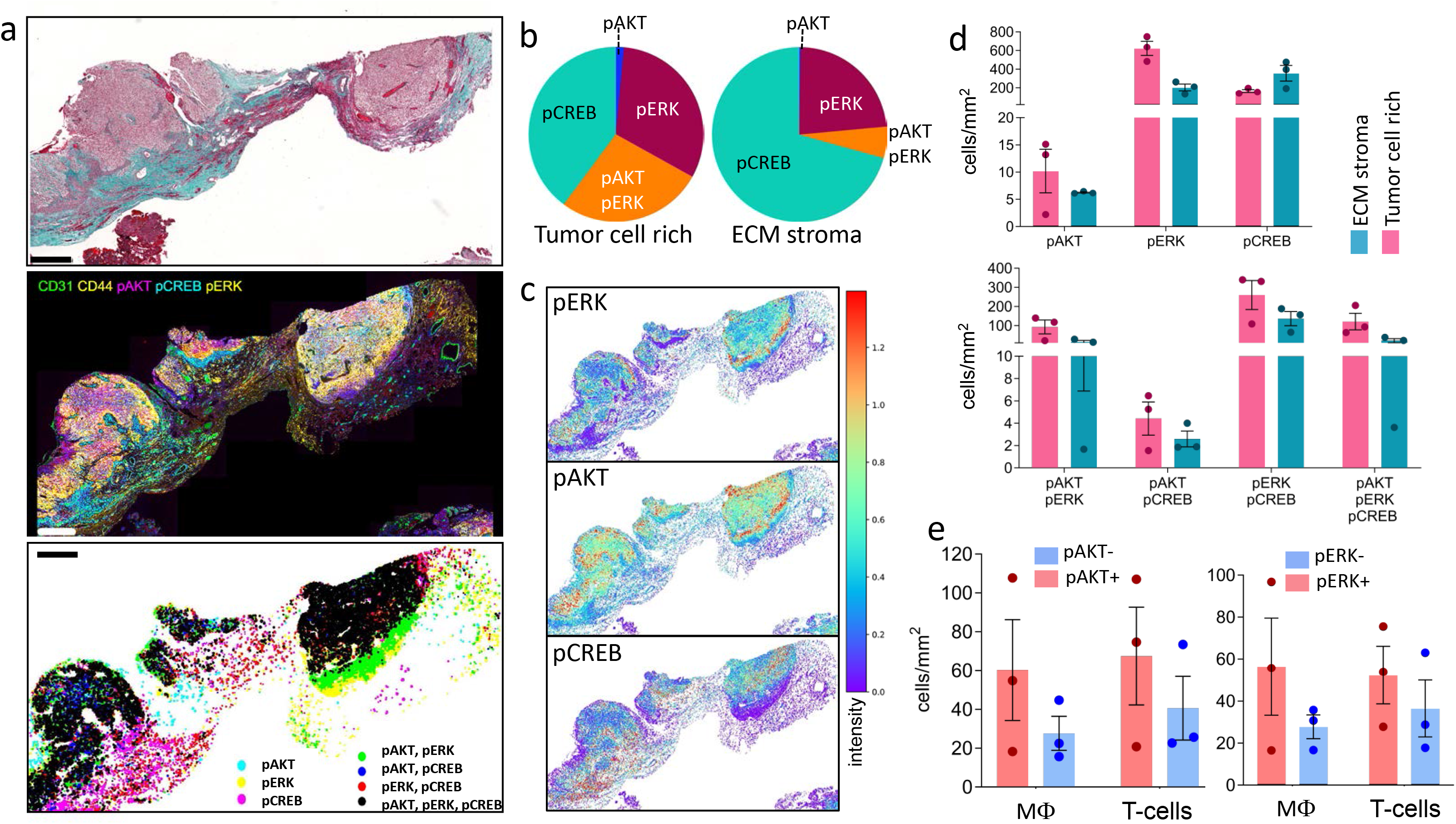
Cell signaling heterogeneity underpins the histopathological and functional heterogeneity and in GBM. **(A)** Representative Masson’s Trichrome staining of a whole GBM tissue, mIHC showing endothelial cells (CD31+), tumor cells (CD44+), PI3K signaling (pAKT+), MAPK signaling (pERK+) and CREB activation (pCREB+) and spatial plot of single or combined signaling positive cells in the same tumor tissue. Scale bars are 100μm. **(B)** Proportion of total pCREB+ cells that are only pCREB positive, and pAKT or pERK positive, and double pAKT and pERK positive. **(C)** Signal intensity mapping for pAKT, pERK and pCREB. **(D)** Cell density of single pAKT, pERK and pCREB positive cells, and all respective combinations is shown as the number of cells per tumor cell rich, and ECM-stroma area analyzed (mm^2^) within whole tumor sections (*n*=3). **e**, Cell densities of macrophages (CD68+) and T-cells (CD3+) are shown as the number of cells per PI3K or MAPK signaling-rich area analyzed (mm^2^) within the whole tumor sections (*n*=3). Error bars are +/- SEM. Statistical significance was determined by Wilcoxon matched-pairs signed rank test and significance is represented by *(p < 0.05), **(p<0.01), ***(p<0.001) and ****(p<0.0001).

Microglia, dendritic cells and T-cells showed a similar even distribution across ECM-stroma and tumor cell rich regions. Considering possible sample bias using small (1.5mm diameter) tissue cores and to further explore intratumoral immune cell distribution, we examined at least four whole biopsy (>10mm^2^) tissue specimens. In most large tissue samples, more immune cells were seen in the ECM-stroma, compared to tumor cell rich regions (Figure 6C).

To determine ECM protein composition, immunohistochemistry was performed to investigate ECM proteins previously reported to be enriched in human brain tissue (Novak and Kaye 2000). Collagen type I-α1 (COL1) was enriched around blood vessels; collagen type IV-α1 (COL4) was enriched in the ECM-stroma, including cells with myofibroblast-like phenotype; fibronectin (FN1) was also enriched in the ECM-stroma and in long fibrillar structures interconnecting stroma-rich regions (Figure S5A). Smooth muscle actin (SMA) exhibited high expression in the ECM-stroma in zones adjacent to tumor cell rich regions and around blood vessels in the ECM-stroma. Vimentin was enriched in the ECM-stroma and associated with vascular rich clusters and was expressed at a lower level in tumor cell rich regions. The expression of tumor growth factor 1 (TGFB) was enriched in the ECM-stroma, with lower expression observed in some tumor cells.

By merging two independent GBM single-cell RNA-sequence data sets derived from 27,589 GBM cells from 12 GBM tumors (Darmanis et al. 2017; Yuan et al. 2018), we investigated the cellular sources of the ECM associated factors, collagen type I and IV, fibronectin, vimentin and smooth muscle actin mRNA expression. *COL1A1* and *COL4A1* expression is most abundant in vascular cells (Figure S5B). Most fibronectin (*FN1*) and vimentin (*VIM*) expressing cells are vascular cells, but myeloid (macrophages and microglia) and glial (astrocyte-like) cells also express high levels of both. Vimentin is also highly expressed by tumor (neoplastic) cells. Vascular cells and neoplastic cells exhibited the highest levels of smooth muscle actin (*ACTA2*) expression. Interrogation of TCGA-LGG-GBM gene expression data and matched patient survival data (Ceccarelli et al. 2016) demonstrates that high mRNA expression of ECM proteins investigated, correlates with poor patient survival (Figure S5C).

### Cell signaling heterogeneity underpins the histopathological and functional heterogeneity and in GBM

We previously reported that cells in GBM tissue exhibit heterogeneous regional cell signaling activation (P. M. Daniel, Filiz, Tymms, et al. 2018). We examined PI3K, MAPK and CREB cell signaling activation in ECM-stroma and tumor cell rich regions, using phospho-AKT (pAKT) as a PI3K activation biomarker, phospho-ERK1/2 (pERK1/2) to determine MAPK activation and pCREB as a biomarker of transcriptional activation, downstream of PI3K and MAPK signaling. Using CD31 to identify vascular tissue/endothelial cells and CD44 to label tumor cells, we identified regional correlation of cell signaling activity in vascular rich stromal regions and tumor cell rich regions. Tumor cell rich tissue regions exhibited a higher proportion of pAKT and pERK1/2 expression, while pCREB expression was more prominent in the ECM-stroma (Figure 7A,B and Figure S6). pAKT and pERK co-expression was enriched in tumor cell rich regions, as was pAKT and pCREB co-expression. Tumor cell dense regions expressed all three (pAKT, pERK, pCREB) signaling biomarkers (Figure 7A,C and Figure S6), while tumor cell dense zones at the ECM-stroma interface exhibited intense pAKT and pERK co-expression (Figure 7C), but lacked pCREB expression. To investigate whether co-expression of pAKT and pERK in tumor cells at the ECM-stroma interface could define an invasive phenotype, neurospheres derived from a brain tumor mouse model cultured in a collagen-rich matrix exhibited strong co-expression of pAKT and pERK in cells invading into the matrix (Figure S7). In highly vascularized (CD31+ rich) ECM-stroma, pCREB and pERK were co-expressed. To determine if there was a correlation between immune cell distribution and cell signaling niches, we measured macrophage and T-cell density in pAKT+ and pERK+ regions, in three different GBM tissues. ECM-stroma exhibited an increased but not significantly different macrophage and T-cell cell density.

## Discussion

Using multiplex immunohistochemistry, we examined patient glioma tissue heterogeneity by measuring cell composition, spatial localization of stromal cells, cell signal activation and tissue ECM remodeling. There was a grade-specific increase in immune cell infiltration, with the infiltration of specific cell types at different stages of glioma development.

T-cell infiltration occurred late in glioma carcinogenesis, evidenced by the presence of T-cells in most grade IV gliomas and little or no detectable infiltration in grade I, II and III tissues. The observations are consistent with previous studies, limited to the analysis of grade II and IV (Weenink et al. 2019). Of the T-cells investigated, Tregs were the rarest cell type, consistent with recent studies (Klemm et al. 2020), suggesting that Tregs may not be key to the establishment of an immunosuppressive tumor immune microenvironment in glioma. Macrophages dominate the glioma tumor microenvironment in all tumor grades, appearing earlier (lower grades), compared to all other immune cell types examined, and macrophages also infiltrate deeper into tumor cell rich regions than other infiltrating immune cells. Previous work using GBM single-cell sequence analysis showed that macrophages are equally distributed across the tumor core and tumor periphery (Darmanis et al. 2017). Immune cell composition in LGG and HGG determined by the tissue analysis data was similar to that predicted by computational deconvolution of bulk RNA-sequencing data. However, our data suggests that there are more infiltrating CD8+ T-cells, compared to CD4+ T-cells, which may reflect the transcriptomic similarity of these T-cell subsets, and a limitation of gene signature-based cell-type prediction.

Although T-cells were present in GBM tissue, most T-cells were localized within perivascular immune cell nests. GBM tumors also showed the presence of tertiary lymphoid structures composed of B-cell, T-cells and macrophages. A recent report showed that the presence of tertiary lymphoid structures in glioma correlates with an enhanced T-cell tumor infiltration, but a reduced T-cell response (van Hooren et al. 2021). Other studies have also reported similar perivascular enriched T-cell localization (Klemm et al. 2020; Weenink et al. 2019), (Robinson et al. 2020). The restricted T-cell localization is likely due to a complex interplay between GBM endothelial cell derived expression of T-cell chemoattractant chemokines such as CXCL12 (Rao et al. 2012), or expression of immunosuppressive cytokines including TGFB1 and IL10 (Cui et al. 2020). Aside from immunosuppressive cytokines and chemokines, our data suggests that immune cells delivered to the tumor via blood vessels face a significant physical barrier due to pathologically high levels of ECM protein deposition around the blood vessels. This may have implications for CART cell-based therapies targeting brain cancer, since efficient delivery of CART cells to brain tumor cells will be impeded by the extensive perivascular ECM and the immunosuppressive tumor microenvironment. This implies that strategies will need to be engineered into CART cell design to overcome these barriers for brain cancer therapy.

Apart from an increased immune cell infiltration in HGG compared to LGG, we observed a grade-dependent increase in the number and proportion of activated immune cells, determined by the expression of phosphorylated CREB. To our knowledge this is the first study using a cell signaling and transcription factor biomarker to identify tumor immune cell activation, in situ. Several studies have relied on computational methods using gene expression signatures to interrogate tumor-derived transcriptomic data to measure stimulus-specific T-cell and dendritic cell activation (K. Yan et al. 2021; Basit et al. 2021; Sun et al. 2021). Immune cell activation in the context of cell signaling pathway activation in the GBM tumor microenvironment highlights the multiple regulatory levels regulating the tumor microenvironment and considerations in the applicability of pathway-specific inhibitors in glioma therapy.

Genomic and transcriptomic studies show that the PI3K and MAPK pathways are critical cell signaling pathways in GBM cell biology (Pearson and Regad 2017; P. M. Daniel, Filiz, Brown, et al. 2018; Fung et al. 2019). Our data identifies heterogeneous cell signaling activity across the GBM tumor microenvironment which correlates with histopathological hallmarks, including tumor cell density, angiogenesis and zones where tumor cell rich regions interface with the ECM-stroma (Figure S6). Tumor cell rich regions exhibit activation of PI3K, MAPK and their downstream transcription factor, CREB, suggesting that these pathways and the CREB transcriptional program are required to sustain cell proliferation, cell invasion and metabolism. We previously showed that in an HGG mouse model in which tumors are initiated by constitutive activation of the PI3K pathway in neural progenitor cells, MAPK and CREB are coactivated, and that targeted CREB deletion in the tumor cells results in slower tumor growth, suggesting that pathway crosstalk and downstream transcriptional programs involving these factors are important for glioma tumorigenesis (P. M. Daniel, Filiz, Brown, et al. 2018). Here, we show that coactivation of the PI3K and MAPK pathways, in the absence of CREB activation, occurs at the stromal-tumor cell rich interface, suggesting that the cells at this interface are less proliferative. In breast cancer, tumor cells at the stromal-tumor interface exhibit no or low proliferative activity, epithelial-mesenchymal transition and express invasive biomarkers (Provenzano et al. 2006; Shen et al. 2014). In an in vitro model, invading brain tumor cells exhibited strong and selective co-activation of PI3K and MAPK signaling in the cells invading into a collagen-rich matrix (Figure S7). PI3K and MAPK pathway coactivation, driven by growth factor receptor signaling, has been shown to be critical for efficient cancer cell invasion(Rambur et al. 2020). In GBM, we also observed high levels of CD44 and CD44v6 expression at the ECM-tumor interface (Figure S4 and data not shown). CD44v6 expression regulates tumor cell invasion and metastasis (Afify, Purnell, and Nguyen 2009; Ma, Dong, and Chang 2019) and is associated with glioma malignancy (Bar et al. 2014), suggesting that activation of both the PI3K and MAPK signaling pathways are required for efficient tumor cell invasion. The lack of pCREB expression we report in GBM, is consistent with the “go or grow” principle, that cell motility and proliferation is typically mutually exclusive (Xie, Mittal, and Berens 2014). Overall, the cell signaling heterogeneity observed in the ECM-stroma and tumor cell rich regions in GBM suggests that both cellular and noncellular components of the tumor microenvironment influence tumor cell signaling. Moreover, ECM-stromal regions enriched in blood vessels showed CREB activation in the absence of PI3K and MAPK signal activation suggesting that CREB may not be required for endothelial cell proliferation. Recognizing the activation status of immune cell types or immune cell subsets will be critical to understanding the tumor-specific functions and ultimately to the specific targeting of the tumor immune microenvironment. Furthermore, identifying the immune cell behaviors associated with pCREB expression, will be critical to understanding specific tumor immune cell functions. An unexpected finding was that ECM deposition in GBM, in patients over 40 years old, was higher in males, compared to females. As glioma typically occurs in older patients and that disease incidence is higher in men, one of the differences contributing to this sex difference may be the higher ECM protein expression in men. Fibrosis-associated disease involving many organs is higher in men and thought to be due to hormone-dependent mechanisms regulating chronic inflammation (Garate-Carrillo et al. 2020). The concept that cancers mirror chronic inflammatory conditions and aberrant wound healing has gained traction recently (MacCarthy-Morrogh and Martin 2020). The identification of collagen deposition, expression of ECM proteins, angiogenesis associated with ECM-stroma and immune cell infiltration in glioma and GBM, as presented in our study, is consistent with the concept that gliomas also share cellular and molecular mechanisms consistent with chronic inflammatory diseases.

The view that the GBM tumor microenvironment is heterogeneous and that the underlying cellular, genetic and molecular factors, establish a biologically complex system which is a barrier to deciphering therapeutically actionable mechanisms. The multilayered tissue analysis approach used in our study provides a greater understanding of the glioma tumor microenvironment, providing spatial context for the deeper understanding of the biological communication networks regulating glioma biology. Moreover, we suggest that suitably engineered macrophages are likely to provide more effective cell-based immunotherapies for glioma and GBM, compared to T-cell based therapies, a concept recently proposed by Joyce and colleagues (Kowal, Kornete, and Joyce 2019). We also suggest the use of simple and inexpensive staining methods to identify collagen deposition in GBM, applied to brain tumor pathology examination, will provide for a more robust diagnosis, especially in distinguishing between grade III and IV glioma. Furthermore, our data suggests that the non-cellular tumor microenvironment in GBM could be targeted. Recent evidence in rat GBM models, suggests that temozolomide alters ECM remodeling in tumors, to generate a pro-tumorigenic microenvironment (Tsidulko et al. 2018). Using anti-fibrotic drugs to disrupt tumor ECM, in combination with standard therapies, could slow the progression of LGG to HGG, and significantly enhance patient survival.

## Methods

### Glioma tissue & Human Ethics

Glioma tissue specimens were from various sources and were all FFPE. Glioma and GBM tissue microarrays were from US Biomax (BS17017b, GL861 and GL806f) and from Lifespan Biosciences (LS-SBRNC14, LS-SBRNC22, and LS-SBRN1). GBM tissue was also sourced from the Department of Anatomical Pathology, Royal Melbourne Hospital. Human ethics approval for tissue use was covered by project application 1853511 and was approved by the Medicine and Dentistry Human Ethics Sub-Committee, The University of Melbourne.

### Multiplex immunohistochemistry, image acquisition and cell phenotyping

Multiplex IHC was performed using an Opal 7-Color IHC Kit (Akoya Biosciences) on a Bond RX automated stainer (Leica Biosystems). All primary antibodies were diluted in the Opal Blocking/Antibody Diluent. A pan immune antibody panel and a T-cell antibody panel were used to investigate immune cell localization in the GBM tumor microenvironment. The pan immune panel included CD68 (Abcam ab955, 1/100), pCREB (Abcam ab32096), TMEM119 (Abcam 185333, 1/1000), CD11c (Abcam ab52632, 1/1000) and CD3 (Abcam ab16669, 1/150). The T-cell subset antibody panel included CD3 (Abcam ab16669, 1/150), CD4 (Abcam ab, 1/250), CD8 (Abcam ab4055, 1/100), FOXP3 (Abcam ab22510, 1/100), PD-1 (Bio SB, 1/100) and pCREB (Abcam ab32096, 1/500). Furthermore, mIHC was also used to identify activation of signaling pathways and angiogenesis using a panel dedicated for different signaling markers, which include pERK (Cell Signaling Technology #9102, 1:200), pAKT (Cell Signaling Technology #4060, 1:100), pCREB (Abcam #ab32096, 1/500), CD44 (Cell Signaling Technology #ab157107, 1:200), CD31 (Dako #M0823, 1:50).

4µm formalin-fixed paraffin-embedded (FFPE) tissue sections were baked at 60°C and deparaffinized in xylene and rehydrated in a serial dilution of ethanol. All slides were then subjected to heat-induced antigen retrieval prior to incubation with 3% hydrogen peroxide. Primary antibody was then added to tissue and incubated at room temperature for 60 minutes, followed by incubation with Opal Polymer HRP Ms + Rb for 30 minutes. Immunofluorescent signals were visualized using Opal fluorophores (Opal 520, 540, 570, 620, 650 and 690), diluted at 1:150 in Plus Automation Amplification Diluent. Serial multiplexing was performed by repeating antigen retrieval, primary and secondary antibody incubation, and Opal polymer visualization until all six markers are added. GBM sections were then counterstained with DAPI and coverslipped using ProLong Glass Antifade Mountant.

### Masson’s Trichrome staining

Masson’s Trichrome staining was performed as previously described (Burkholder 1974). FFPE tissue sections were deparaffinized in xylene and rehydrated through decreasing grades of ethanol. Slides were incubated with Carazzi haematoxylin followed by an incubation with a 1% Briebrich scarlet and 1% acid fuchsin solution. Tissue sections were decolorized with 1% phosphotungstic acid and subsequently incubated with 1% light green prior to a brief rinse with two changes of absolute ethanol and xylene, followed by coverslipping.

### Microscopy and image analysis

Multiplex IHC images were acquired using the Vectra 3.0.5 Multispectral Imaging Platform (Perkin Elmer, USA) at 40x magnification. Spectral deconvolution was then performed using inForm 2.4.4 (Perkin Elmer, USA). High-resolution multispectral images were then fused and analyzed using the software Halo (Indica Labs, USA). Masson’s Trichome stained images were acquired using a Vectra 3.0.5 Multispectral Imaging Platform (Perkin Elmer, USA) at 10x magnification. Image analysis was carried out using Halo, where cell phenotyping was performed using the analysis algorithm Highplex FL 3.0.3, according to the cell phenotyping matrix (Supplementary Table 1). Cell phenotyping involved adjusting the nuclear detection sensitivity, establishing the minimum intensity threshold for positive detection, and defining biomarker localization as nuclear, cytoplasmic, or membranous. To determine the level of collagen/ECM stroma, the Tissue Classifier add-on in Halo was used. Tissue Classifier uses a machine learning algorithm to identify tissue types based on user input software training.

### Spatial Analysis

Spatial analysis was performed on Halo to assess macrophage and T-cell tumor infiltration by annotating perivascular areas on mIHC images and annotating ECM-stroma and tumor cell rich regions from Masson’s Trichrome stained tissue. Cell localization was identified by measuring the distance of plotted cells to the nearest perivascular location using the Infiltration Analysis Module in Halo. Immune cells and PI3K or MAPK signaling positive cells in tumor cell rich regions and in the ECM-stroma were enumerated based on regions identified using Masson’s Trichrome staining, cell phenotypes and thresholding for signal detection on Halo. To compare the number of macrophages and T-cells in PI3K and MAPK signaling regions, CD68 and CD3 positive cells were analyzed in pAKT or pERK1/2 signaling dense regions annotated on serial tissue sections using Halo. Signaling spatial and intensity plots were graphed using the Matplotlib (version 3.4.3)(Hunter 2007) package in Python, based on object data from image analysis in Halo.

### Statistical analysis

Statistical analyses included a Kruskal-Wallis test, followed by Dunn’s test for multiple comparisons, a two-tailed unpaired Student’s t-test and Wilcoxon matched-pairs signed rank test with GraphPad Prism v.9 software, as indicated in each figure legend. Statistical significance is represented by *(p<0.05), **(p<0.01), ***(p<0.001) and ****(p<0.0001).

## Acknowledgments

This work was supported by the CASS Foundation Australia (grants 6236 and 7941), Department of Surgery (RMH) and School of Biomedical Sciences translational grant. We thank Metta Jana and Ian Birchall for advice on tissue preparation, histology and staining. Marlene Hao for reagents and discussions.

## Author contributions

M.D. and S.S.W. performed the majority of experimental work, data analysis and wrote parts of the manuscript. L.F. and L.C. performed immunohistochemical analysis experiments. Y.F. and S.M. performed the computational analysis experiments. P.N., P.D. and R.R. provided expertise on multiplex immunohistochemical analysis optimization experiments. F.M. provided expertise on tumor immunology and antibody panel selection. S.S. provided expertise for glioma tissue selection and co-conceived the project. T.M. is the principal investigator who conceived the project, supervised research and wrote and edited the manuscript.

## SUPPLEMENTARY DATA FIGURE LEGENDS

**Figure S1.**
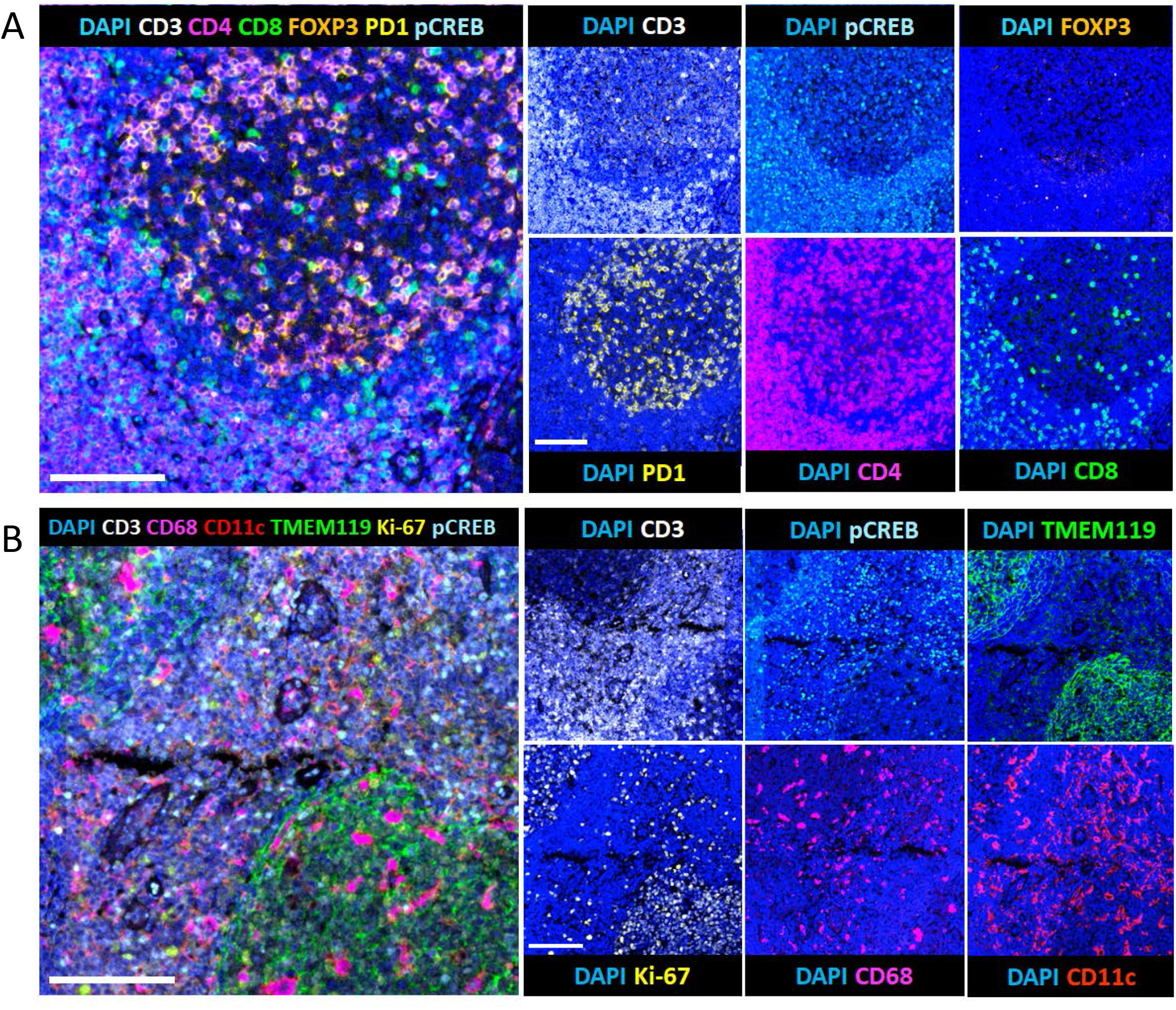
mIHC antibody panel validation. Human tonsil tissue sections (4μm thick) were stained with antibodies and conditions shown in Table S1. Multispectral images were captured using a Vectra 3.0.5 microscope, unmixed on inForm 2.4.4 software, and analyzed using HALO software, as described in the Methods. Scale bars are 100μm.

**Figure S2.**
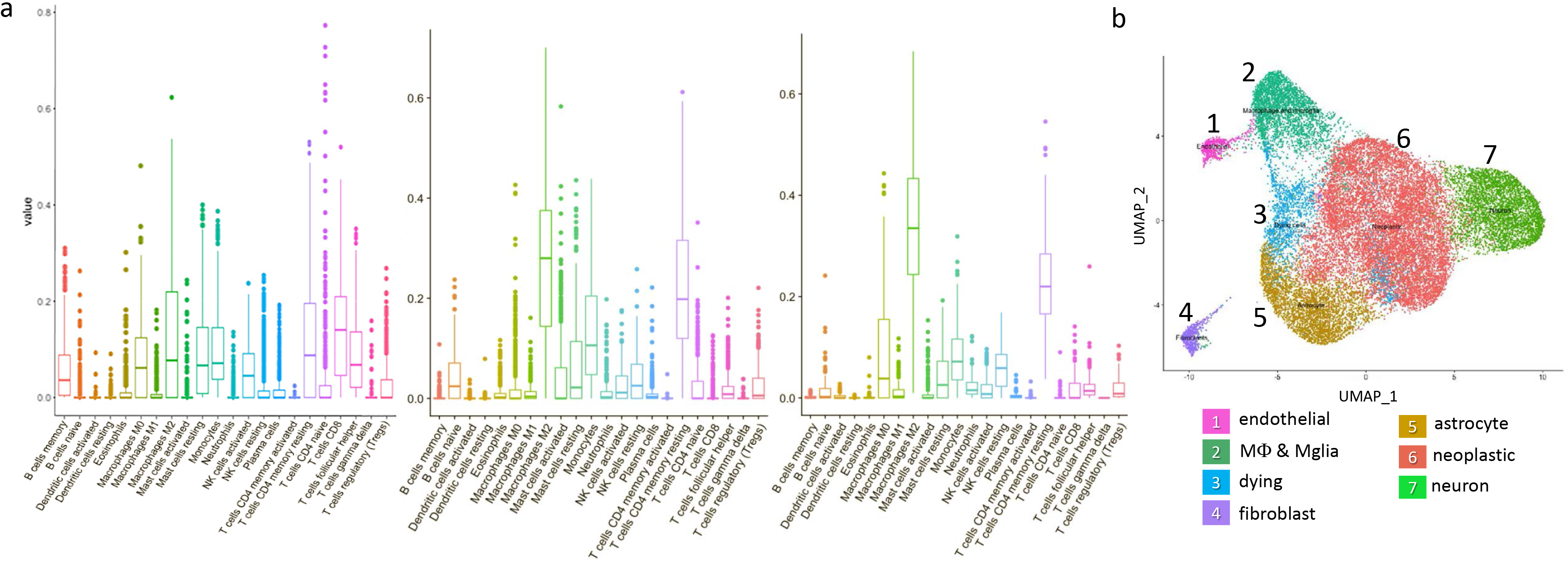
Glioma cell composition predicted by computational deconvolution of gene expression data. **(A)** Immune cell composition in non-tumor brain (NT), low-grade glioma (LGG) and glioblastoma (GBM) estimated by computational deconvolution using CIBERSORT (Chen et al. 2018). NT brain expression data was derived from prenatal brain tissue (Miller et al. 2014) and glioma gene expression data sets were from LGG and GBM tissue (Ceccarelli et al. 2016). **(B)** Cell identification in glioblastoma was determined by uniform manifold approximation and projection (UMAP), using single-cell RNA-sequencing data merged from two studies which analyzed a total of 27,589 GBM cells, isolated from 12 GBM tumors (Darmanis et al. 2017; Yuan et al. 2018).

**Figure S3.**
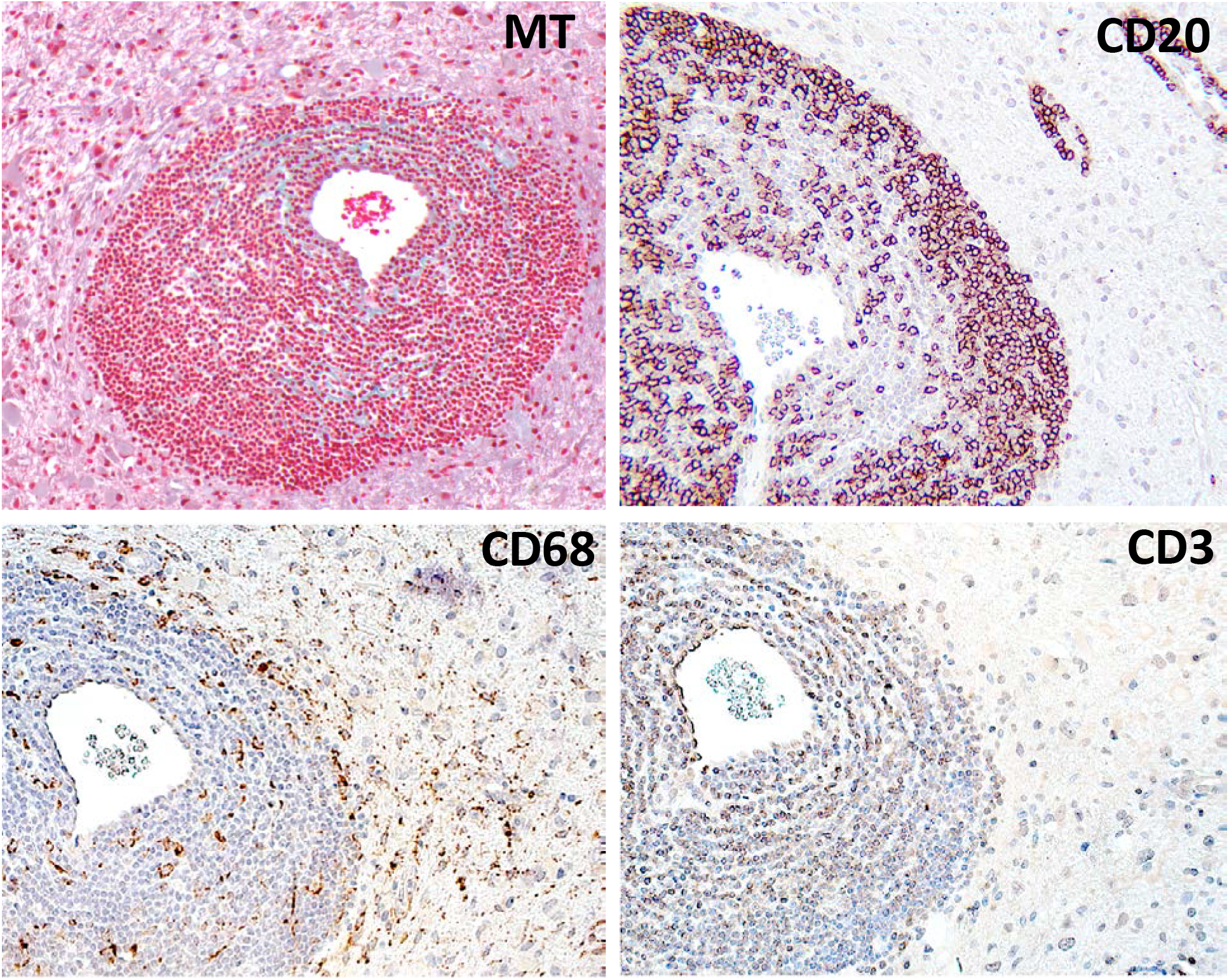
Vascular-associated lymphoid structures (VALS) in glioblastoma. VALS, visualized by Masson’s trichrome stain (MT), exhibited dense concentric arrangement of immune cells, with intervening collagen fibers (light green). Immune cell identity was determined by immunohistochemistry. VALS were composed of B-cells (CD20), macrophages (CD68) and T-cells (CD3). Scale bars are 50μm.

**Figure S4.**
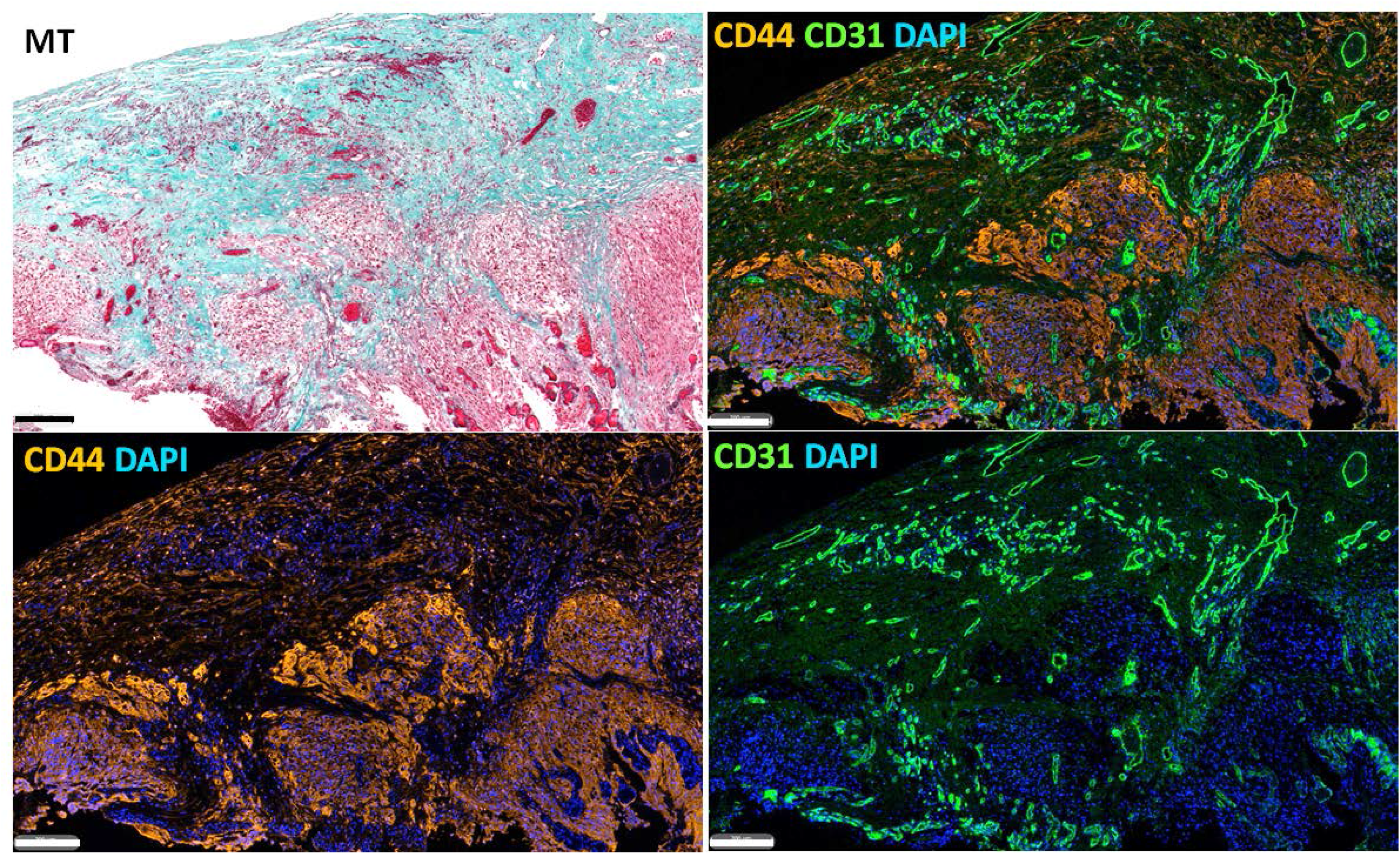
ECM-stroma associated angiogenesis. Masson’s trichrome (MT) stain was used to distinguish ECM-stroma, characterized by collagen deposition (green). Tumor cell rich regions are red. Immunofluorescence was used to identify tumor cells (CD44, orange) and endothelial cells (CD31, green). Scale bars are 200μm.

**Figure S5.**
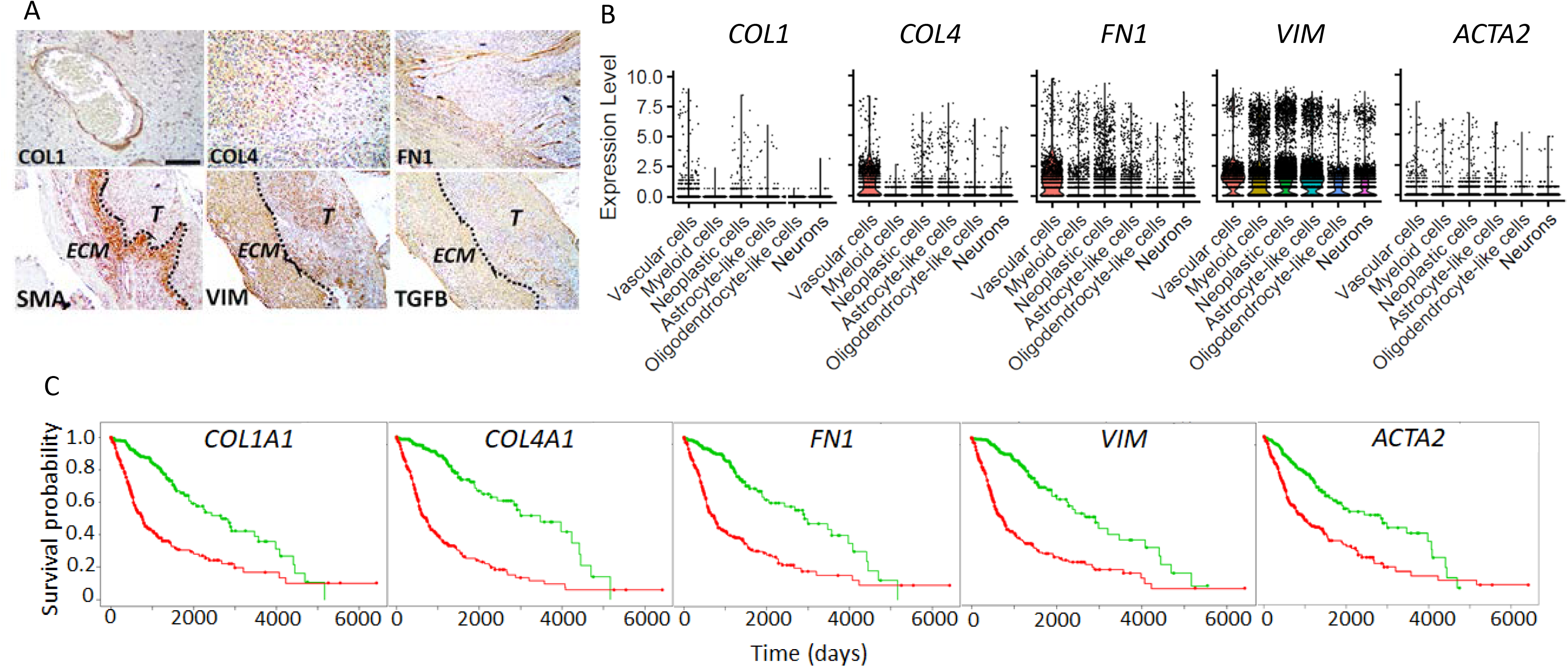
ECM-associated and myofibroblast protein and mRNA expression in GBM stroma. **(A)** Collagen I (COL1) protein was enriched around blood vessels; collagen IV (COL4) protein was expressed in angiogenic-rich ECM-stroma, fibronectin protein was enriched in the ECM-stroma and ECM-associated fibers; smooth muscle actin (SMA) protein was enriched in the ECM-stroma, at the ECM-tumor interface. Vimentin (VIM) and tumor growth factor-β (TGFB) were enriched in the ECM-stroma and tumor cell clusters. T denotes the tumor-cell rich region and ECM denotes the ECM stromal region; the dotted line marks the ECM-tumor cell-rich interface. **(B)** Normalized gene expression for ECM associated mRNAs is shown for different tumor and brain cell types. Normalized expression was determined from single-cell RNA-sequencing data merged from two studies, which analyzed a total of 27,589 GBM cells, isolated from 12 GBM tumors(Darmanis et al. 2017; Yuan et al. 2018). **(C)** Kaplan-Meier plots showing LGG and GBM patient survival and correlation with ECM-associated mRNA expression. Correlation was determined using TCGA LGG-GBM data(Ceccarelli et al. 2016).

**Figure S6.**
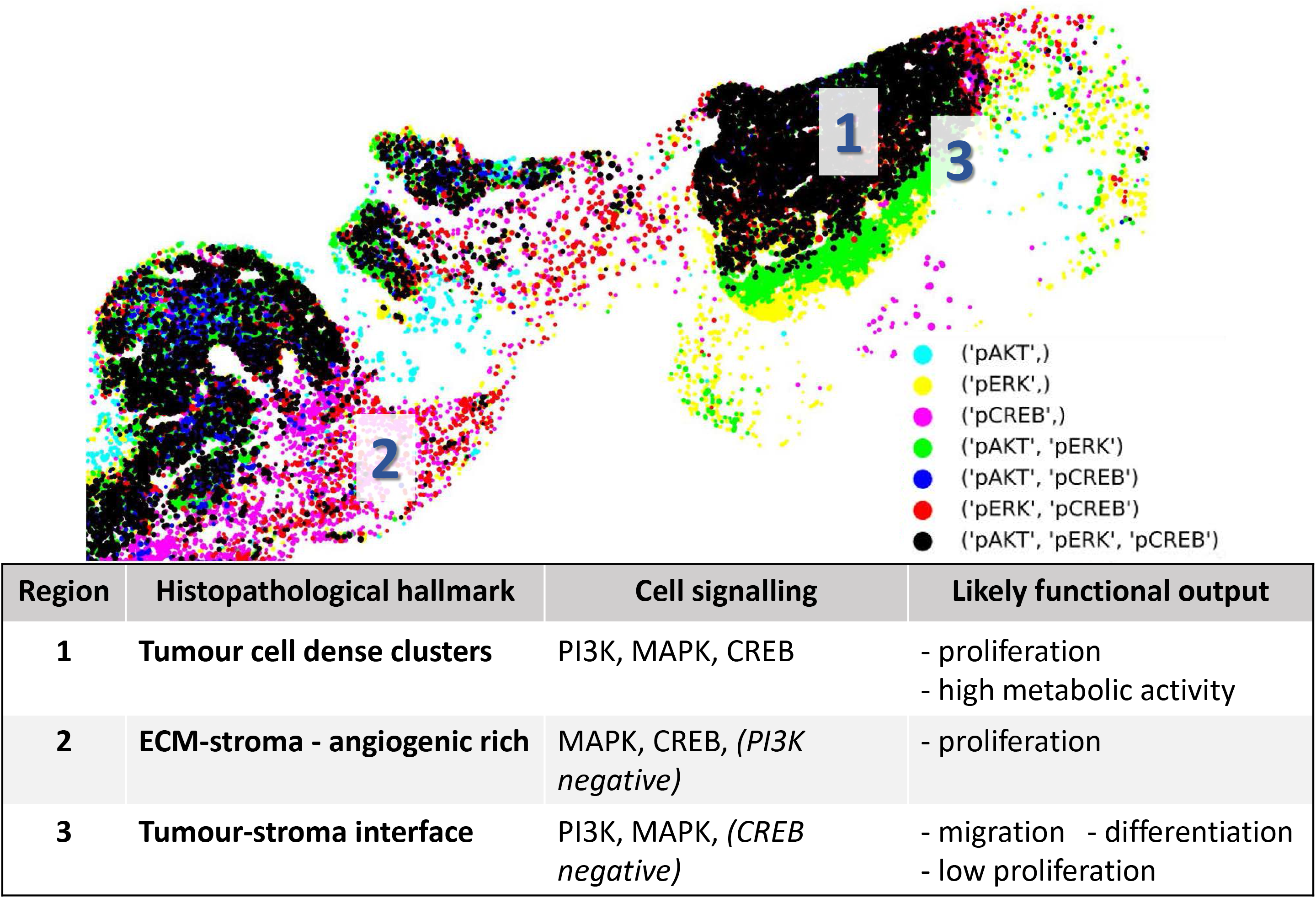
Spatial analysis of cell signaling activity in GBM defines cellular function. Signaling spatial and intensity plots were plotted using the Matplotlib, using object identifier data from image analysis in Halo, as described in the Methods. Cell signaling profiles are indicated in the color key at the bottom right. Three regions (1-3) are shown and the corresponding dominant cell signaling activity and likely functional output is listed in the table. Functions were predicted based on our prior data (P. M. Daniel, Filiz, Brown, et al. 2018; P. M. Daniel, Filiz, Tymms, et al. 2018; P. Daniel et al. 2014).

**Figure S7.**
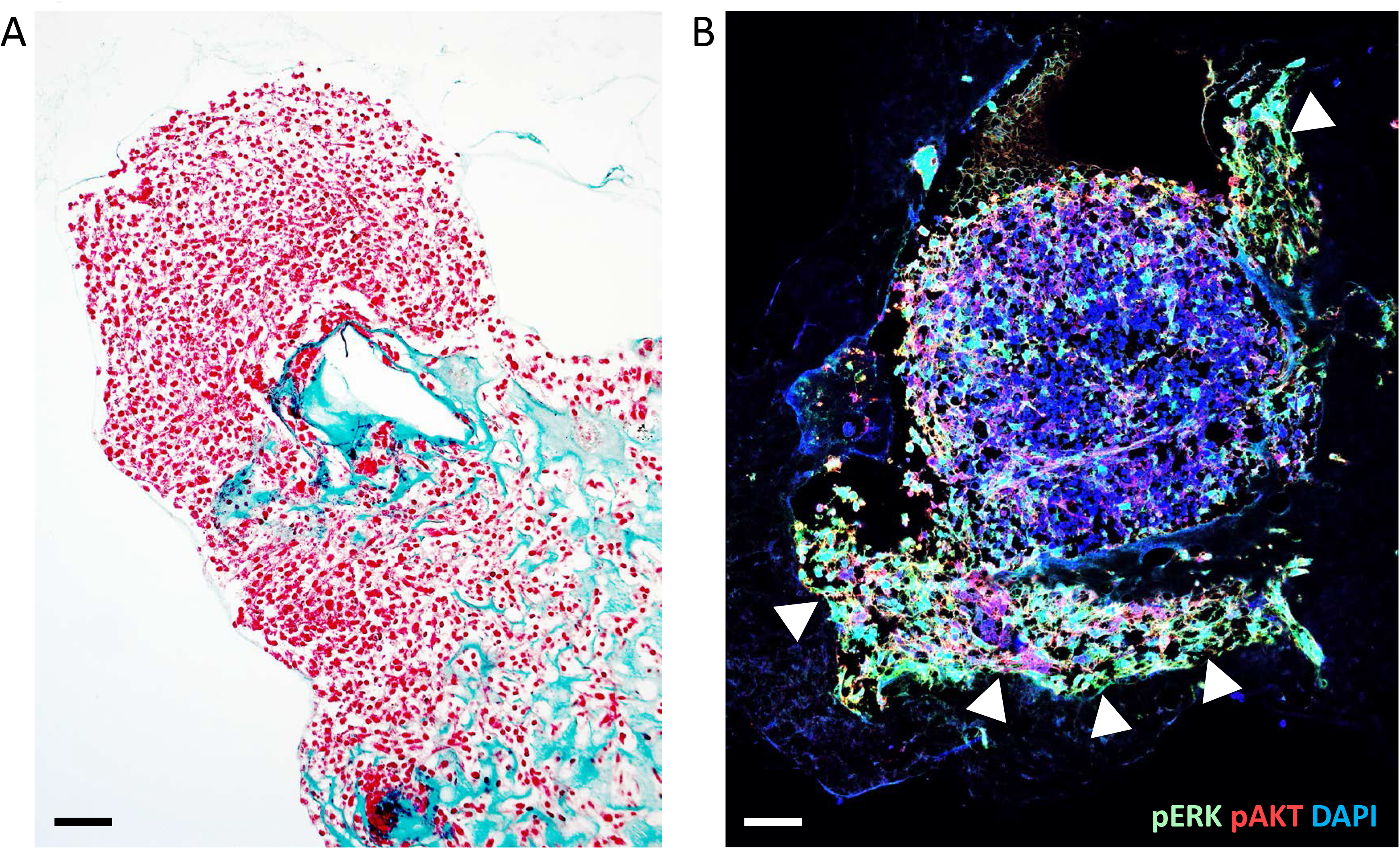
Co-activation of the PI3K and MAPK pathways in invasive brain cancer cells. **(A)** Mouse brain cancer cells, cultured as neurospheres, were placed into a collagen-rich matrix and incubated at 37°C for 24 hours. **(B)** Cells invading (arrow heads) into the matrix exhibit intense PI3K (pAKT, green) and MAPK (pERK, red) co-expression.

**Supplementary Table 1.** Cell biomarker phenotyping matrix used to identify cell types in glioma tissue

| Cell type | Biomarkers |
| --- | --- |
| **Tumour cell** | DAPI<br>(AND) pCREB (OR) Ki-67<br>(NOT) CD3, TMEM119, CD68, CD11c |
| **Dendritic cell** | DAPI<br>(AND) CD11c<br>(NOT) CD3, CD68, TMEM119 |
| **Microglia** | DAPI<br>TMEM119<br>(NOT) CD3 |
| **Macrophage** | DAPI<br>(AND) CD68<br>(NOT) CD3, TMEM119 |
| **T-cell** | DAPI<br>(AND) CD3<br>(NOT) CD11c, CD68, TMEM119 |
| **Treg** | DAPI<br>(AND) CD3<br>(AND) FOXP3<br>(NOT) CD11c, CD68, TMEM119 |
| **other T-cells** | DAPI<br>(AND) CD3<br>(AND)/(NOT) CD4 (OR) CD8 (AND/NOT) PD-1 |

**Supp Table 2.** Immune cell composition in glioma. Cell density (cells/mm2) and the % proportion of each cell type relative to all DAPI^+^ cells across the tissue segment examined.

|  | **GRADE** | **NT** | **I** | **II** | **III** | **IV** |
| --- | --- | --- | --- | --- | --- | --- |
| **Microglia** | cells/mm<sup>2</sup> | 31.3 | 20.6 | 31 | 87 | 32.8 |
|  | % of cells | 2.35 | 0.83 | 1.2 | 2.4 | 1.1 |
| **Macrophages** | cells/mm<sup>2</sup> | 1.6 | 70 | 56 | 55 | 133 |
|  | % of cells | 0.1 | 3.5 | 2.1 | 1.2 | 3.7 |
| **Dendritic cells** | cells/mm<sup>2</sup> | <0.1 | 23.4 | 17.5 | 5.8 | 19 |
|  | % of cells | <0.1 | 1.7 | 0.6 | 0.2 | 0.6 |
| **T-cells** | cells/mm<sup>2</sup> | <0.1 | 0.9 | 4.8 | 2 | 14 |
|  | % of cells | <0.1 | <0.1 | 0.2 | 0.1 | 0.4 |

**Supp Table 3.** T-cell subtype composition in glioma. Cell density (cells/mm2) and the % proportion of each cell type relative to all T-cells across the tissue segment examined.

| T-cell | GRADE | NT | I | II | III | IV |
| --- | --- | --- | --- | --- | --- | --- |
| CD3+CD4+ | cells/mm<sup>2</sup> | <0.1 | <0.1 | <0.1 | 2.5 | 1.5 |
|  | % of cells | <0.1 | <0.1 | <0.1 | 19.4 | 8.1 |
| CD3+CD8+ | cells/mm<sup>2</sup> | <0.1 | 1 | 0.5 | 6.1 | 4.8 |
|  | % of cells | <0.1 | 32 | 15.6 | 37.6 | 28.6 |
| CD4+PD1+ | cells/mm<sup>2</sup> | <0.1 | <0.1 | <0.1 | <0.1 | <0.1 |
|  | % of cells | <0.1 | <0.1 | <0.1 | <0.1 | <0.1 |
| CD8+PD1+ | cells/mm<sup>2</sup> | <0.1 | <0.1 | <0.1 | <0.1 | <0.1 |
|  | % of cells | <0.1 | <0.1 | <0.1 | <0.1 | <0.1 |

